# Stable transplantation of human mitochondrial DNA by high-throughput, pressurized mitochondrial delivery

**DOI:** 10.1101/2020.09.15.298174

**Authors:** Alexander J. Sercel, Alexander N. Patananan, Tianxing Man, Ting-Hsiang Wu, Amy K. Yu, Garret W. Guyot, Shahrooz Rabizadeh, Kayvan R. Niazi, Pei-Yu Chiou, Michael A. Teitell

## Abstract

Generating mammalian cells with specific mtDNA-nDNA combinations is desirable but difficult to achieve and would be enabling for studies of mitochondria-nucleus communication and coordination in controlling cell fates and functions. We developed ‘MitoPunch’, a force-actuated mitochondrial transfer device, to deliver isolated mitochondria into numerous target mammalian cells simultaneousl*y*. MitoPunch and MitoCeption, an alternative force-based mitochondrial transfer approach, both yield stable isolated mitochondrial recipient (SIMR) cells that permanently retain exogenous mtDNA, whereas coincubation of mitochondria with cells does not yield SIMR cells. Although a typical MitoPunch or MitoCeption delivery results in dozens of immortalized SIMR clones with restored oxidative phosphorylation, only MitoPunch can produce replication-limited, non-immortal human SIMR clones. The MitoPunch device is versatile, inexpensive to assemble, and easy to use for engineering mtDNA-nDNA combinations to enable fundamental studies and potential translational applications.

## INTRODUCTION

Mitochondrial (mtDNA) and nuclear (nDNA) genome coordination regulates metabolism, epigenome modifications, and other processes vital for mammalian cell survival and activity (Patananan et al., 2018; Ryan & Hoogenraad, 2007; Singh et al., 2017). Together, these genomes encode >1,100 mitochondrial proteins, with only 13 essential electron transport chain (ETC) proteins encoded within the mtDNA (Calvo & Mootha, 2010). Mutations in mtDNA can impair the ETC by altering nDNA co-evolved ETC complex protein interactions, causing defective respiration and debilitating diseases (Greaves et al., 2012). Techniques that exchange non-native for resident mtDNAs could enable studies of mtDNA-nDNA interactions and replace deleterious mtDNAs within cells with therapeutic potential (Patananan et al., 2016).

Our current inability to edit mtDNA sequences is a roadblock for many studies and potential applications. For example, endonucleases targeted to the mitochondrion inefficiently eliminate and cannot alter mtDNA sequences (Bacman et al., 2018). Mitochondrial transfer between cells *in vitro* and *in vivo* provides a potential path forward for transplanting existing mtDNA sequences, however, the mechanisms controlling such transfers remain unknown (Dong et al., 2017; Torralba et al., 2016). Methods that deliver mitochondria into mtDNA-deficient (so-called ‘ρ0’) cells include membrane disruption (King & Attardi, 1988; Wu et al., 2016) or fusion with enucleated cytoplasts (Wilkins et al., 2014). However, these methods are typically laborious, low-throughput, or depend on cancerous, immortal recipient cells lacking physiologic mitochondrial activity. An interesting recent study did report one desired mtDNA-nDNA clone and 11 false-positive clones using cybrid fusion with replication-limited cells, an achievement hampered by a low generation rate with unknown reproducibility or generalizability (Wong et al., 2017). Thus, a higher throughput, reproducible, and versatile mtDNA transfer approach to generate multiple desired ‘stable isolated mitochondrial recipient’ (SIMR) clones in replication-limited cells remains essential for statistically valid studies and potential translation of mitochondrial transplantation.

## RESULTS

### MitoPunch mechanism uses fluid pressure to disrupt the plasma membrane

We developed ‘MitoPunch’ as a simple, high-throughput mitochondrial transfer device consisting of a lower polydimethylsiloxane (PDMS) reservoir loaded with a suspension of isolated mitochondria, covered by a polyethylene terephthalate (PET) filter seeded with ~2 x 10^5^ adherent cells. Upon actuation, a mechanical plunger deforms the PDMS from below, which, by numerical simulation, generates pressure up to 28 kPa inside the PDMS chamber, propelling the suspension through numerous 3μm pores in the PET filter. This pressure cuts the plasma membrane of recipient cells sitting atop the pores and delivers mitochondria into the cytoplasm of the cut cells (Figures 1A and 1B). To assess performance, we compared MitoPunch to mitochondrial coincubation (Kitani et al., 2014) and to MitoCeption, a method that also uses pressure to deliver mitochondria into mammalian cells. In MitoCeption a 1500 x g centripetal force exerts a pressure of ~1.6 Pa to associate mitochondria with recipient cell membranes (Figures 1C and 1D) (Caicedo et al., 2015).

**Figure 1.**
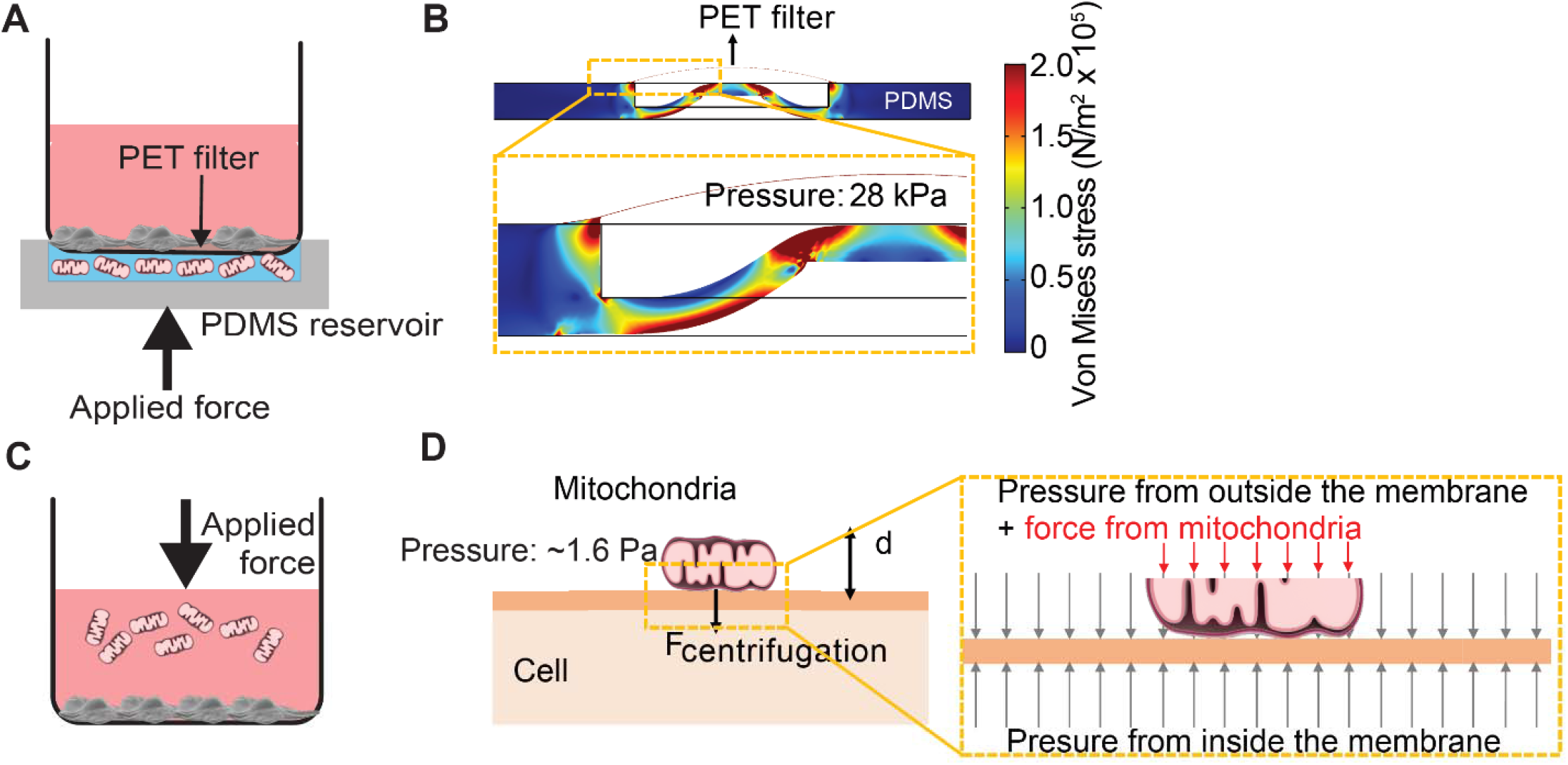
Pressure simulations of mitochondrial transfer tools. **(A)** Schematic of MitoPunch apparatus. Recipient cells (1 x 10^5^) were seeded on a porous PET membrane ~24 h before delivery. A freshly isolated suspension of mitochondria in 1x PBS, pH 7.4, is loaded into the PDMS chamber and the filter insert is sealed over the PDMS before activation of the mechanical plunger to pressurize the apparatus and deliver the mitochondrial suspension into recipient cells. **(B)** Numerical simulation showing the pressure inside the PDMS chamber reaching 28 kPa with piston activation. **(C)** Schematic of MitoCeption technique. **(D)** MitoCeption pressure model.

### Mitochondrial delivery into transformed and primary cells

We isolated and transferred dsRed-labeled mitochondria from ~1.5 x 10^7^ HEK293T cells (Miyata et al., 2014) into ~2 x 10^5^ 143BTK− ρ0 osteosarcoma cells and replication-limited BJ ρ0 foreskin fibroblasts and measured the fraction of recipient cells positive for dsRed fluorescence. For 143BTK− ρ0 cells at ~2 h post-delivery, flow cytometry showed that MitoPunch yielded the lowest number of dsRed-positive cells compared to coincubation or MitoCeption. Similarly, for BJ ρ0 recipient cells, MitoPunch yielded the lowest number of dsRed-positive cells compared to coincubation or MitoCeption, with all values reduced relative to 143BTK− ρ0 recipients (Figure 2A). These data suggest that the method of delivery and target cell type affect mitochondrial transfer efficiency for each method.

**Figure 2.**
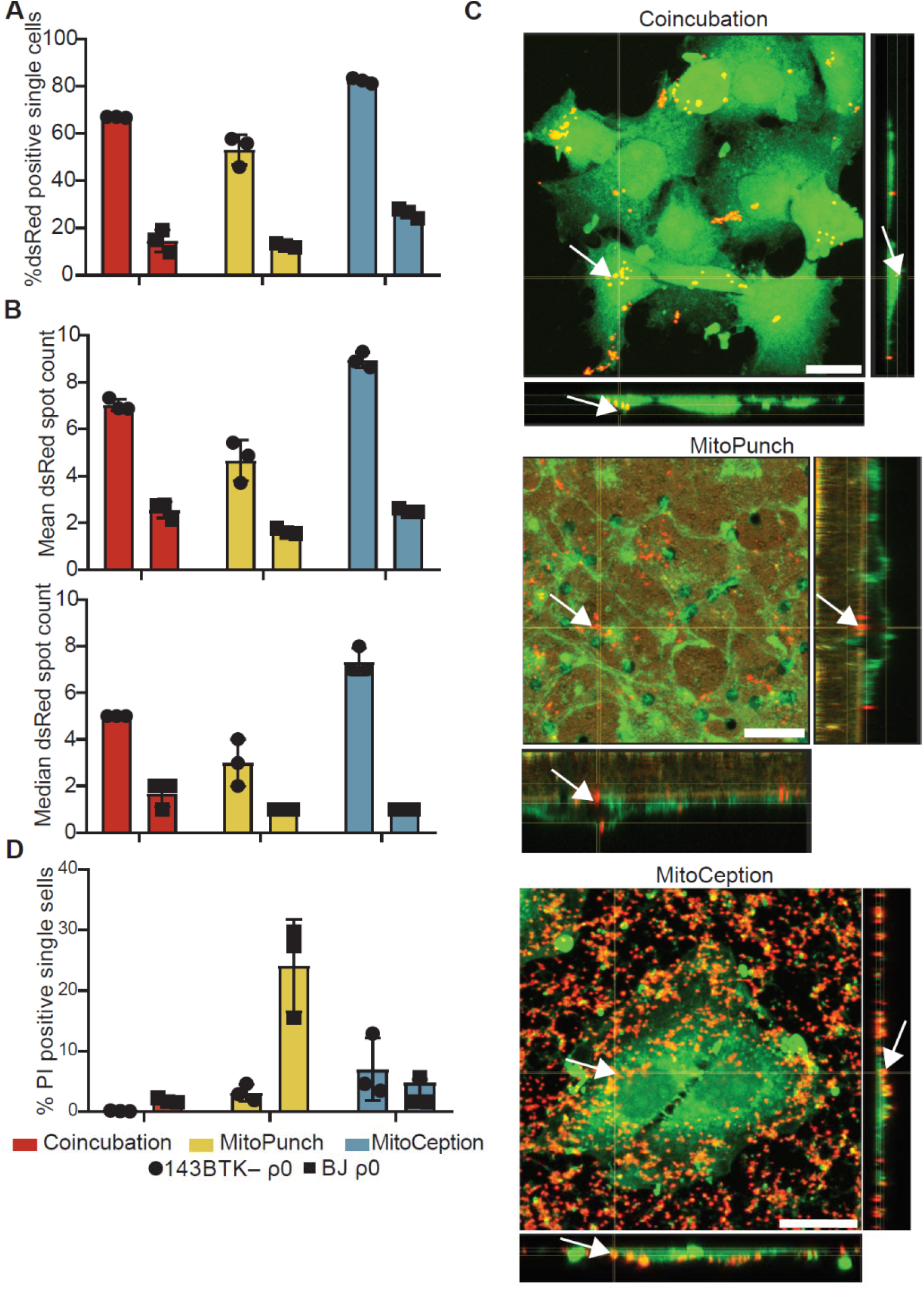
MitoPunch delivers isolated mitochondria into recipient cells. **(A)** Quantification of flow cytometry results measuring dsRed mitochondria within 143BTK− ρ0 and BJ ρ0 single cells following mitochondrial transfer. **(B)** Mean and median dsRed spot count quantification of ImageStream data. **(C)** Sequential z-stacks of confocal microscopy of isolated HEK293T dsRed mitochondria delivered into 143BTK− ρ0 cells from the basal to apical face of the cell in descending order. Arrows indicate representative mitochondria interacting with recipient cells. Red is dsRed mitochondria and green in coincubation and MitoCeption is CellMask Green plasma membrane stain and wheat germ agglutinin plasma membrane stain in MitoPunch. Scale bars indicate 15 μm. **(D)** Quantification of flow cytometry measurements of fluorescence in 143BTK− ρ0 and BJ ρ0 single cells following propidium iodide transfer by coincubation. Error bars repre-

We quantified the number of discreet dsRed-spots in each cell ~2 h following delivery using ImageStreamx MarkII imaging flow cytometry (George et al., 2004) (Figures 2A and S1A). ImageStream spot count analysis of 143BTK− ρ0 recipient cells showed MitoPunch delivered a lower mean and median number of dsRed spots per cell than coincubation or MitoCeption. MitoPunch transfers into BJ ρ0 recipient cells yielded fewer mean spots/cell compared to coincubation and MitoCeption with an equivalent median number of spots/cell for MitoPunch and MitoCeption and slightly more for coincubation. Next, we used confocal microscopy to observe HEK293T dsRed mitochondrial fluorescence 15 min post-transfer in the 143BTK− ρ0 recipient, selected for its robust mitochondrial acquisition. We found that MitoPunch localized mitochondria to pores in the filter insert and the cytoplasm of cells, whereas MitoCeption uniformly coated recipient cells with mitochondria (Figure 2C). While both methods transfer mitochondria to recipient cells, MitoPunch delivers mitochondria to filter pore-exposed sections of the plasma membrane, compared to a diffuse membrane association pattern seen with MitoCeption.

To complement fluorescence microscopy and flow cytometry, we imaged mitochondrial recipient cells by scanning electron microscopy (SEM) 15 min after transfer (Figure S1B). SEM analysis of the filter and basal membranes shows that MitoPunch generates pores of ~1 μm in recipient cells. In contrast, MitoCeption coats recipient cells with mitochondria and no observable pores form in recipient cell membranes. We next delivered the membrane impermeant dye propidium iodide (PI) by coincubation, MitoPunch, or MitoCeption to measure membrane disruption from delivery and quantified uptake by flow cytometry (Novickij et al., 2017). Delivery into 143BTK− ρ0 cells by MitoPunch and MitoCeption resulted in similar percentages of PI-positive recipient cells, and both were greater than coincubation. Interestingly, BJ ρ0 cells showed comparable fractions of PI-positive cells to the 143BTK− ρ0 after coincubation and MitoCeption. However, MitoPunch yielded a ~5-fold increase in the PI-positive fraction compared to all other conditions (Figure 2D). These data show that MitoPunch and MitoCeption disrupt the plasma membrane of recipient cells for potential mitochondrial transfer, and the degree of disruption is cell type and delivery method dependent.

### Stable retention of transplanted mtDNA

After verifying that MitoPunch and MitoCeption successfully transfer mitochondria to recipient cells, we next tested their SIMR cell generation capability. ρ0 cells cannot synthesize pyrimidines and therefore cannot proliferate or survive without added uridine because of ETC impairment, so we used uridine-free medium as selection to isolate SIMR cells with transplanted mtDNA and restored ETC activity (Gregoire et al., 1984) (Figures 3A and 3B). We transferred HEK293T mitochondria into 143BTK− ρ0 and BJ ρ0 cells by coincubation, MitoPunch, and MitoCeption, cultured recipients in uridine-free media for 7 d, and quantified the resultant number of viable clones by crystal violet staining and manual counting (Figure 3C). Coincubation did not generate SIMR clones in 143BTK− ρ0 cells, in contrast to MitoPunch and MitoCeption, which each generated dozens of clones. BJ ρ0 cells delivered HEK293T mitochondria by coincubation or MitoCeption did not form SIMR clones. MitoPunch generated numerous SIMR clones in both cell types, although fewer BJ ρ0 SIMR clones than in 143BTK− ρ0 cells, whereas MitoCeption only generated clones in 143BTK− ρ0 cells and was unable to form stable clones in replication-limited BJ cells.

**Figure 3.**
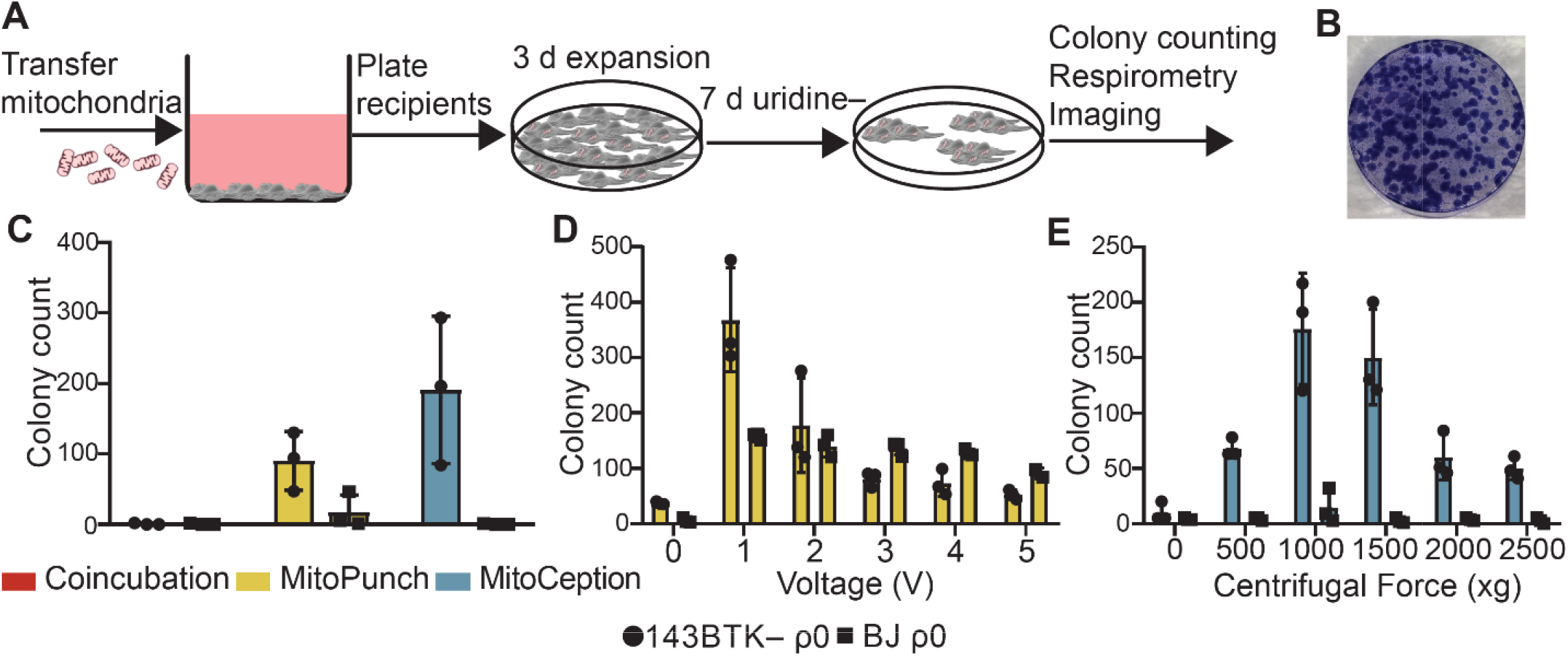
Stable retention of transplanted mtDNAs into transformed and replication-limited cells. **(A)** Workflow for SIMR cell generation by mitochondrial transfer into ρ0 cells. **(B)** Representative crystal violet stained plate following MitoPunch and SIMR cell selection. **(C)** Quantification of crystal violet stained 143BTK− ρ0 and BJ ρ0 SIMR clones. Error bars represent SD of three technical replicates. **(D)** Quantification of crystal violet stained 143BTK− ρ0 and BJ ρ0 SIMR clones formed by MitoPunch actuated with indicated voltages after uridine-free selection. Error bars represent SD of three technical replicates with the exception of BJ ρ0 5 V transfer, which shows two replicates. **(E)** Quantification of crystal violet stained 143BTK− ρ0 and BJ ρ0 SIMR clones formed by MitoCeption with indicated centripetal forces after uridine-free selection. Error bars represent SD of three technical replicates.

We next investigated whether differences in SIMR clone generation between 143BTK− ρ0 and BJ ρ0 cells is driven by sensitivity to differences in delivery pressure. We developed a MitoPunch device with adjustable plunger acceleration modulated by changing the circuit voltage. We achieved maximum 143BTK− ρ0 SIMR clone generation with this tunable MitoPunch at 1 V, with a sharp reduction to background with increasing voltage (Figure 3D). The BJ ρ0 recipient also showed maximal SIMR generation at 1 V, with a shallow decline in SIMR generation efficiency to 5 V. Additionally, MitoPunch deliveries into B16 ρ0 mouse melanoma cells (Tan et al., 2015) yielded maximal SIMR generation at a different voltage than in the human cell lines tested, showing that optimal mitochondrial delivery pressure may be cell type dependent (Figure S2A). We performed a similar force titration with MitoCeption by adjusting the maximum centripetal force. In 143BTK− ρ0 cells, we observed maximum clone generation at 1000 x g and we obtained no BJ ρ0 SIMR clones at any acceleration tested for MitoCeption (Figure 3E). These data suggest that MitoPunch is uniquely able to generate SIMR clones in replication-limited fibroblasts and SIMR generation efficiency depends on delivery pressure.

To enable desirable mtDNA-nDNA clone generation using limited starting material, such as mitochondria from rare cell subpopulations, we determined the minimal biomass of mitochondrial isolate required to generate SIMR clones. We performed coincubation, MitoPunch, and MitoCeption transfers into 143 BTK− ρ0 recipient cells using HEK293T dsRed mitochondrial suspensions of decreasing concentrations. After delivery, we seeded half of each sample for SIMR cell selection in uridine-free medium and analyzed the fraction of cells that received mitochondria in the remaining half by flow cytometry. We observed a similar dose-dependent relationship for MitoPunch and MitoCeption for 0.16 μg, 1.6 μg, and 16 μg total mitochondrial protein suspended in 120 μl of 1x DPBS, pH 7.4 transfer buffer (Figure S2B). The SIMR clone generation efficiency per 1 x 10^4^ dsRed-positive cells from the 16 μg mitochondrial suspension deliveries was greater for MitoPunch than MitoCeption. These results show that although MitoPunch and MitoCeption generate SIMR clones with similar efficiency per μg of mitochondrial isolate, MitoPunch generates >4 times as many stable clones per mitochondrial recipient in a transformed cell type. A significant volume of the original 120 μl mitochondrial suspension remains in the PDMS delivery chamber following MitoPunch, so we tested whether the same sample could be used for serial transfers. We performed 11 sequential deliveries into 143BTK− ρ0 cells using one aliquot of mitochondrial isolate and found maximal SIMR clone generation from the first and second deliveries, after which SIMR cell formation dropped off by the 11^th^ transfer (Figure S2C).

### SIMR cells rescue ρ0 mitochondria network morphology and respiration

Finally, we measured mitochondrial function in SIMR cells by quantifying the rate of oxygen consumption and assessed mitochondrial morphology. We isolated three independent 143BTK− ρ0 SIMR clones generated by MitoPunch or MitoCeption transfer of HEK293T mitochondria and measured each clone’s oxygen consumption rate (OCR) using a Seahorse Extracellular Flux Analyzer mitochondrial stress test (Figure 4A). To determine whether SIMR clone respiration remained stable through time and tissue culture passaging, we grew four clones, two from MitoPunch and two from MitoCeption, through two freeze/thaw cycles and measured their respiration. We found that one MitoCeption clone lost its respiratory capacity and one MitoPunch clone was not viable after freeze down and thaw (data not shown). In the remaining clones, basal and maximal respiration, spare respiratory capacity, and ATP generation remained stable throughout both freeze thaw cycles (Figure 4A). We then imaged the SIMR clones for TOM20 mitochondrial outer membrane protein and double-stranded DNA levels by confocal microscopy (Figures 4B and S3A). The MitoCeption clone that lost respiratory capacity showed a fragmented mitochondrial network with no detectable mtDNA, whereas the other SIMR clones generated by MitoPunch and MitoCeption contained mtDNA with filamentous mitochondrial network morphologies. These data showed that the majority of 143BTK− ρ0 SIMR clones generated by either MitoPunch or MitoCeption have stable levels of mtDNA content and restored respiratory profiles.

**Figure 4.**
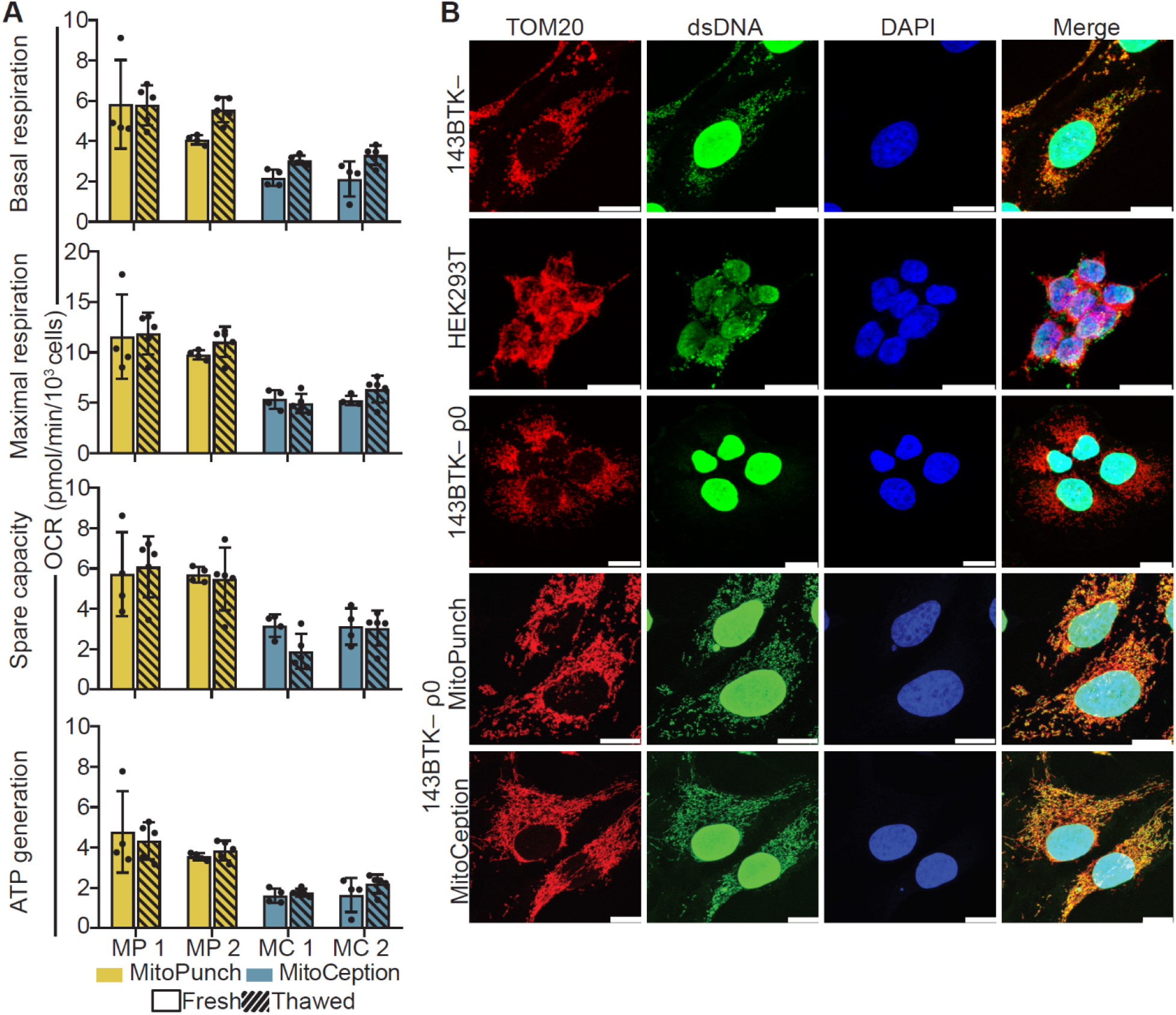
mtDNA transplantation rescues ρ0 mitochondrial phenotype. **(A)** OCR quantification of basal and maximal respiration, spare respiratory capacity, and ATP generation from two independent 143BTK− ρ0 + HEK293T SIMR clones generated by MitoPunch and MitoCeption. Cross-hatched data indicate clones that were frozen and thawed twice each. Error bars represent SD of four technical replicates for fresh measurements and five for thawed SIMR cell measurements. **(B)** Confocal microscopy of representative 143BTK− ρ0 + HEK293T SIMR clones compared to 143BTK− parental, HEK293T dsRed mitochondrial donor, and 143BTK− ρ0 controls. Scale bars indicate 15 μm.

## DISCUSSION

Stability of the mitochondrial genome is essential for studying the long-term effects of mtDNA-nDNA interactions and for potential therapeutic applications of mitochondrial transfer. MitoPunch generates up to hundreds of SIMR clones in both transformed and Hayflick-limited recipient cells by exerting a pressure sufficient to perforate mammalian cell membranes in regions small enough to be repaired within seconds to minutes, which sustains cell viability and resumed cell growth and proliferation (Boye et al., 2017). Studies in our laboratory suggest that the transcriptome and metabolome of replication-limited SIMR clones differs significantly from un-manipulated control clones but can be recovered and reset to un-manipulated control clone levels by cellular reprogramming to induced pluripotent stem cells and subsequent differentiation (Patananan, et al., under review). These results indicate that SIMR clone generation in replication-limited, reprogrammable cells is crucial for studies of mtDNA-nDNA interactions, and that MitoPunch is uniquely capable of efficiently generating enough clones for statistically valid studies in such work.

## MATERIALS AND METHODS

### Cell culture conditions

Human ρ0 cells were grown in DMEM (Fisher Scientific, Waltham, MA, Cat. # MT10013CM) supplemented with 10% FBS, non-essential amino acids (Gibco, Waltham, MA, Cat. #11140-050), GlutaMax (Thermo Fisher Scientific, Waltham, MA, Cat. # 35050-061), penicillin and streptomycin (VWR, Radnor, PA, Cat. # 45000-652), and 50 mg/L uridine (Thermo Fisher Scientific, Cat. # AC140770250). All other human cell lines were grown in DMEM (Fisher Scientific, Cat. # MT10013CM) supplemented with 10% FBS, non-essential amino acids, GlutaMax, and penicillin and streptomycin. B16 ρ0 cells were grown in RPMI (Thermo Fisher Scientific, Cat. # MT-10-040-CM) supplemented with 10% FBS, non-essential amino acids, GlutaMax, penicillin and streptomycin, pyruvate (Corning, Corning, NY, Cat. # 25000CI), and 50 mg/L uridine. L929 cells were grown in RPMI supplemented with 10% FBS, non-essential amino acids, GlutaMax, penicillin and streptomycin, and pyruvate. All mammalian cells were maintained in a humidified incubator maintained at 37 °C and 5% CO_2_. The following cells were used in this study: HEK293T (female), 143BTK− (female), 143BTK− ρ0 (female), BJ (male), BJ ρ0 (male), B16 (male), and L929 (male). We have not formally identified these cell lines, however, we have sequenced their mitochondrial and nuclear DNA for polymorphisms and find unique sequences which we use for genotyping our cultures (unpublished data).

### Mitochondrial isolation

Mitochondria were isolated from ~1.5 x 10^7^ mitochondrial donor cells per mitochondrial transfer using the Qproteome Mitochondrial Isolation Kit (Qiagen, Hilden, Germany, Cat. #37612) according to the manufacturer’s instructions. Isolations were done using a 26G blunt ended needle (VWR, Cat. # 89134-164) attached to a 3 ml syringe (VWR, Cat. # BD309657) to mechanically disrupt mitochondria donor cells and did not include high purity isolation procedures. Isolated mitochondrial pellets were re-suspended in 1x DPBS with calcium and magnesium (Thermo Fisher Scientific, Cat. # 14040133) immediately prior to mitochondrial transfer and kept on ice.

### Mitochondrial coincubation

Mitochondria isolated from HEK293T dsRed cells re-suspended in 1x DPBS with calcium and magnesium at ~1mg total protein/ml were pipetted on top of the culture medium of adherent 143BTK− ρ0 and BJ ρ0 recipient cells and incubated at 37°C and 5% CO_2_ for 2 h.

### MitoPunch

The construction and actuation of the MitoPunch device is described in more detail in Patananan, et al, under review, 2020. MitoPunch uses a solenoid-activated impeller to transfer by force isolated mitochondria in a holding chamber into the cytosol of mammalian cells. A 5V solenoid (Sparkfun, Boulder, CO, Cat. # ROB-11015) is positioned on a platform (Thor Labs, Newton, NJ, Cat. # SM1PL) and mounted on a plate (Thor Labs, Cat. # CP02T). A flexible polydimethylsiloxane (PDMS, 10:1 ratio of Part A base: Part B curing agent) (Fisher Scientific, Cat. #NC9644388) reservoir is situated above the solenoid using optomechanical assembly rods (Thor Labs, Cat. # ER3), an upper plate (Thor Labs, Cat. # CP02), and an aluminum washer (outer diameter, 25 mm; inner diameter, 10 mm). The PDMS reservoir consists of a bottom layer (25 mm diameter, 0.67 mm height) bonded to an upper ring (outer diameter, 25 mm; inner diameter, 10 mm; height, 1.30 mm). This reservoir can contain up to ~120 μl of liquid. 1 x 10^5^ adherent cells are seeded on a membrane with 3 μm pores (Corning, Cat. # 353181) 1 d prior to mitochondrial delivery. 120 μl mitochondrial suspension of ~1mg total protein/ml in DPBS with calcium and magnesium is loaded into the PDMS reservoir. Mitochondrial transfer is performed by securing the seeded membrane to the PDMS reservoir. The solenoid, regulated by the Futurlec mini board (Futurlec, New York, NY, Cat. # MINIPOWER) and powered by a power supply (MEAN WELL, New Teipei City, Teiwan, Cat. # RS-35-12), is activated and strikes the middle of the PDMS chamber, displacing the base layer by ~1.3 mm. This displacement pressurizes the mitochondrial suspension and propels it through the membrane and into the cells. In addition, a prototype based on the same principles with identical delivery procedures as MitoPunch, but with tunable plunger force achieved by varying actuator voltage, was engineered by NanoCav, LLC.

### MitoCeption

As described previously (Caicedo et al., 2015), 1 x 10^5^ cells were seeded in one well of a 6-well dish and incubated at 37°C and 5% CO_2_ overnight. Mitochondrial isolate suspended in 1x DPBS with calcium and magnesium at ~1mg total protein/ml was pipetted into the well and the plate was centrifuged at 1500 x g for 15 min at 4°C. Cells were removed from the centrifuge and incubated for 2 h at 37°C and 5% CO_2_ before being centrifuged a second time at 1500 x g for 15 min at 4°C. Cells were then released from the dish and placed into 10 cm plates for SIMR cell selection or harvested for additional analyses.

The pressure generated by the MitoCeption method was estimated by calculating the force exerted per unit area of the cell membrane during centrifugation. The force induced by the centrifugation of a single mitochondria on the cell membrane was equal to the centripetal force of the mitochondria under the acceleration of 1500 x g minus the buoyancy force,

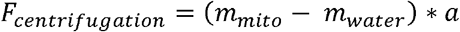
where *m_mito_* and *m_water_* are the mass of mitochondria and water, *a* is the acceleration rate of centrifugation. The equivalent pressure induced by mitochondria centrifugation was approximated by

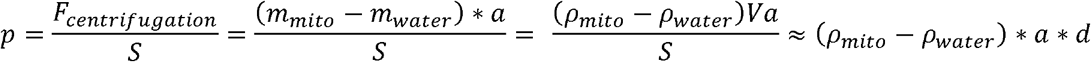
where *ρ_mito_* (1.1 g/cm^3^) and *ρ_water_* (1.0 g/cm^3^) are the density of mitochondria and water, *V* and *S* are the volume and cross-sectional area of mitochondria, and *d* is the thickness of a mitochondrion (~1 μm). Using values for the geometry and properties of a mitochondrion, the pressure induced by MitoCeption centrifugation was ~1.6 Pa.

### Numerical simulation

The finite element method (COMSOL Inc, Burlington, MA, Multiphysics 5.3) was used to simulate the pressure inside the MitoPunch PDMS chamber. We constructed the simulation geometry according to real device dimensions. Piston movement was applied as initial displacement in the y direction. Considering the incompressibility of the aqueous medium inside the PDMS chamber, the volume of the chamber was maintained constant while solving for the stress distribution of all the materials.

### SIMR clone isolation

Mitochondrial recipient and vehicle delivery control 143BTK− ρ0 cells were grown in complete medium supplemented with 50 μg/ml uridine for 3 d following mitochondria or vehicle transfer. After 3 d, the medium was changed to complete medium with 10% dialyzed FBS (Life Technologies, Carlsbad, CA, Cat. # 26400-044) and cells fed daily. After the vehicle delivery control sample died and clones emerged on mitochondrial transfer plates (~7 d in dialyzed FBS complete medium), the medium was changed back to uridine supplemented medium and clones were counted by microscopy or isolated using cloning rings. Mitochondrial recipient and vehicle delivery control BJ ρ0 cells were grown in complete media supplemented with 50 *μ* g/ml uridine for 3 d following mitochondria transfer. After 3 d, the medium was changed to complete medium with 10% dialyzed FBS and cells fed daily. On 5 d post-delivery, cells were shifted to glucose-free, galactose-containing medium (DMEM without glucose, Gibco, Cat. # 11966025) supplemented with 10% dialyzed FBS and 4.5 g/l galactose. After the vehicle delivery control sample died and clones emerged on mitochondrial transfer plates (~36 h in dialyzed FBS galactose medium), the medium was changed to uridine-free medium and colonies were counted by microscopy or isolated using cloning rings.

### ImageStream flow cytometry

Mitochondria were transferred to recipient cells, harvested, and collected in 1.5 ml tubes. Samples were centrifuged 5 min at 1000 x g, supernatant was aspirated, and cells were washed 3x with 0.5 ml 1x PBS, pH 7.4. The PBS was aspirated and cells fixed in 100 μl freshly diluted 4% paraformaldehyde (Thermo Fisher Scientific, Cat. # 28906) for 15 min on ice. Fixative was diluted with 1 ml of 1x PBS, pH 7.4, and 5% FBS and centrifuged for 10 min at 500 x g. Supernatant was removed and cells re-suspended in 1x PBS, pH 7.4, with 5% FBS, images acquired on an ImageStream MarkII platform, and analyzed using the IDEAS 6.2 software package (Luminex, Austin, TX).

### Confocal microscopy

Cells (1 x 10^5^) were plated in 6-well dishes with 2 ml of media on glass coverslips (Zeiss, Oberkochen, Germany, Cat. # 474030-9000) for ~24 h prior to sample preparation. For imaging MitoPunch samples, 1 x 10^5^ cells were plated onto filter inserts 24 h prior to sample preparation. Immediately prior to mitochondrial transfer, coincubation and MitoCeption samples were stained with 1X CellMask Green PM (Molecular Probes, Eugene, OR, Cat. # C37608) diluted in warm medium for 10 min and washed twice in PBS, and MitoPunch samples were stained with 5 μg/mL Alexa Fluor 488 conjugated Wheat Germ Agglutinin (Invitrogen, Carlsbad, CA, Cat. # W11261) diluted in warm media for 10 min and washed twice in PBS. Following delivery, culture medium was removed and 1 ml freshly diluted 4% paraformaldehyde in 1x PBS, pH 7.4, was pipetted onto samples and incubated for 15 min at RT. Paraformaldehyde was aspirated and samples were washed 3x with 1x PBS, pH 7.4. Samples were further washed with PBS 3x with 5 min RT incubation per wash. MitoPunch filters were removed from the plastic insert. Samples were treated with a 10 min RT incubation in 0.1% Triton-X 100 (Sigma, St. Louis, MO, Cat. # X100) to permeabilize cells. Permeabilized samples were washed 3x with 1x PBS and then incubated for 1 h at RT with 2% bovine serum albumin (BSA) dissolved in 1x PBS blocking buffer. Blocking buffer was aspirated and cells incubated for 1 h at RT with a 1:1,000 dilution of primary antibodies in 2% BSA blocking buffer against dsDNA (Abcam, Cambridge, United Kingdom, Cat. # ab27156) and TOM20 protein (Abcam, Cat. # ab78547), and then washed 3x with 5 min RT incubation with 1x PBS. Cells were then incubated with secondary antibodies (Invitrogen, Cat. # A31573 and A21202) diluted 1:100 in 2% BSA blocking buffer protected from light for 1 h at RT. After incubation with secondary antibodies, samples were washed 3x with 5 min 1x PBS incubations and mounted on microscope slides.

To mount, samples were removed from the 6-well dish and rinsed by dipping in deionized water, dried with a Kimwipe, and mounted using ProLong Gold Antifade Mountant with DAPI (Invitrogen, Cat. # P3691) or ProLong Glass Antifade Antifade Mountant with NucBlue™ Stain (Thermo Fisher Scientific, Cat # P36985) on microscope slides (VWR, Cat. # 48311-601). Samples were dried at RT protected from light for 48 h prior to confocal imaging with a Leica SP8 confocal microscope.

### Scanning electron microscopy

Cells (1 x 10^5^) were plated in 6-well dishes with 2 ml of media on glass coverslips ~24 h prior to sample preparation. For imaging MitoPunch samples, 1 x 10^5^ cells were plated onto filter inserts ~24 h prior to sample preparation. Culture medium was removed, and 1 ml freshly diluted 4% paraformaldehyde in 1x PBS was pipetted onto samples and incubated for 15 min at RT. Paraformaldehyde was aspirated and samples were washed 3x with 1x PBS and further washed with 1x PBS 3x with 5 min RT incubation per wash. MitoPunch filters were removed from the plastic insert. Samples were dehydrated by serial 10 min incubations in EtOH:H_2_O solutions of 50%, 60%, 70%, 80%, 90% and 100% EtOH. The samples were immersed into hexamethyldisilazane (HMDS) for 7 min three times and left at RT overnight to air-dry before being coated with 30 nm thick gold/palladium (Au/Pd) before scanning electron microscope (SEM) using a sputter coater (Denton Desk II, Denton Vacuum, Moorestown, NJ, USA).

### Crystal violet staining and clone counting

Media was aspirated from 10 cm plates before fixation with 1 ml freshly diluted 4% paraformaldehyde in 1x PBS for 15 min at RT. Fixative was removed and 1 ml 0.5% w/v crystal violet solution (Thermo Fisher Scientific, Cat. # C581-25) dissolved in 20% methanol in water was applied to each plate and incubated for 30 min at RT. Crystal violet was removed and plates were washed 2x with deionized water before drying overnight at RT and visual quantification.

### Oxygen consumption rate measurements

Oxygen consumption rate (OCR) measurements used a Seahorse XF96 Extracellular Flux Analyzer (Agilent, Santa Clara, CA). 2 x 10^4^ cells were seeded into each well of a V3 96-well plate (Agilent, Cat. # 101085-004) and cultured 24 h before measuring OCR. The Agilent Seahorse mitochondrial stress test was used to quantify OCR for basal respiration and respiration following the sequential addition of mitochondrial inhibitors oligomycin, carbonyl cyanide-p-trifluoromethoxyphenylhydrazone (FCCP), and rotenone.

### Propidium iodide staining, delivery, and flow cytometry

Cells (1 x 10^5^) were plated for delivery and incubated overnight. Media was changed to FluorBrite DMEM media (ThermoFisher Scientific, Cat. # A1896701) with 3 μM propidium iodide (Thermo Fisher Scientific, Cat. # P1304MP) immediately before transfer. MitoCeption and coincubation were carried out as described above, and MitoPunch was performed with PI FluorBrite medium loaded into the PDMS reservoir and incubated for 15 min at 37°C and 5% CO_2_. All samples were washed with 1x PBS and released from culture dishes using Accutase (Thermo Fisher Scientific, Cat # A1110501). Samples were collected in flow cytometry tubes and centrifuged 5 min at 500 x g. Samples were washed with 1x PBS with 5% FBS three times and analyzed on a BD Fortessa flow cytometer.

### Quantification and statistical analysis

All information pertaining to experimental replication is found in the figure legends. All error bars in this manuscript represent standard deviation of three replicates unless otherwise specified in the figure legend.

## ACKNOWLEDGEMENTS

A.J.S. is currently supported by the NIH National Research Service Award fellowship T32CA009120 and previously T32GM007185. A.N.P. is supported by the NIH (T32CA009120) and American Heart Association (18POST34080342). A.K.Y. is supported by the NIH (T32GM008042). G.W.G. is supported by the J.W. and Nellie McDowell Scholarship Fund and by the Boyer Scholarship Fund. P.-Y.C. was supported by the National Science Foundation (CBET 1404080), by the NIH (R01GM114188), and by the AFOSR (FA9550-15-1-0406). M.A.T. is supported by the Air Force Office of Scientific Research (FA9550-15-1-0406), the NIH (R01GM114188, R01GM073981, R01CA185189, R21CA227480, and P30CA016042), and by CIRM (RT3-07678). We thank Rebeca Acin-Perez, Linsey Stiles, and Orian Shirihai of the UCLA Metabolomics Core for help with Seahorse XF Analyzer assays. We thank Zoran Galic, Alejandro Garcia, and Salem Haile of the UCLA Jonsson Comprehensive Cancer Center Flow Cytometry Core Laboratory and Felecia Codrea, Jessica Scholes, and Jeffrey Calimlim of the UCLA Broad Stem Cell Research Center Flow Cytometry Core for assistance with cellular analysis. We thank Laurent Bentolila and Matthew J. Schibler of the UCLA Advanced Light Microscopy/Spectroscopy core for assistance with confocal microscopy. We also thank Emma Dawson, Lynnea Waters, Natasha Carlson, and Christopher Sercel for critical feedback and assistance with this manuscript.

---

**Supplemental Figure 1.**
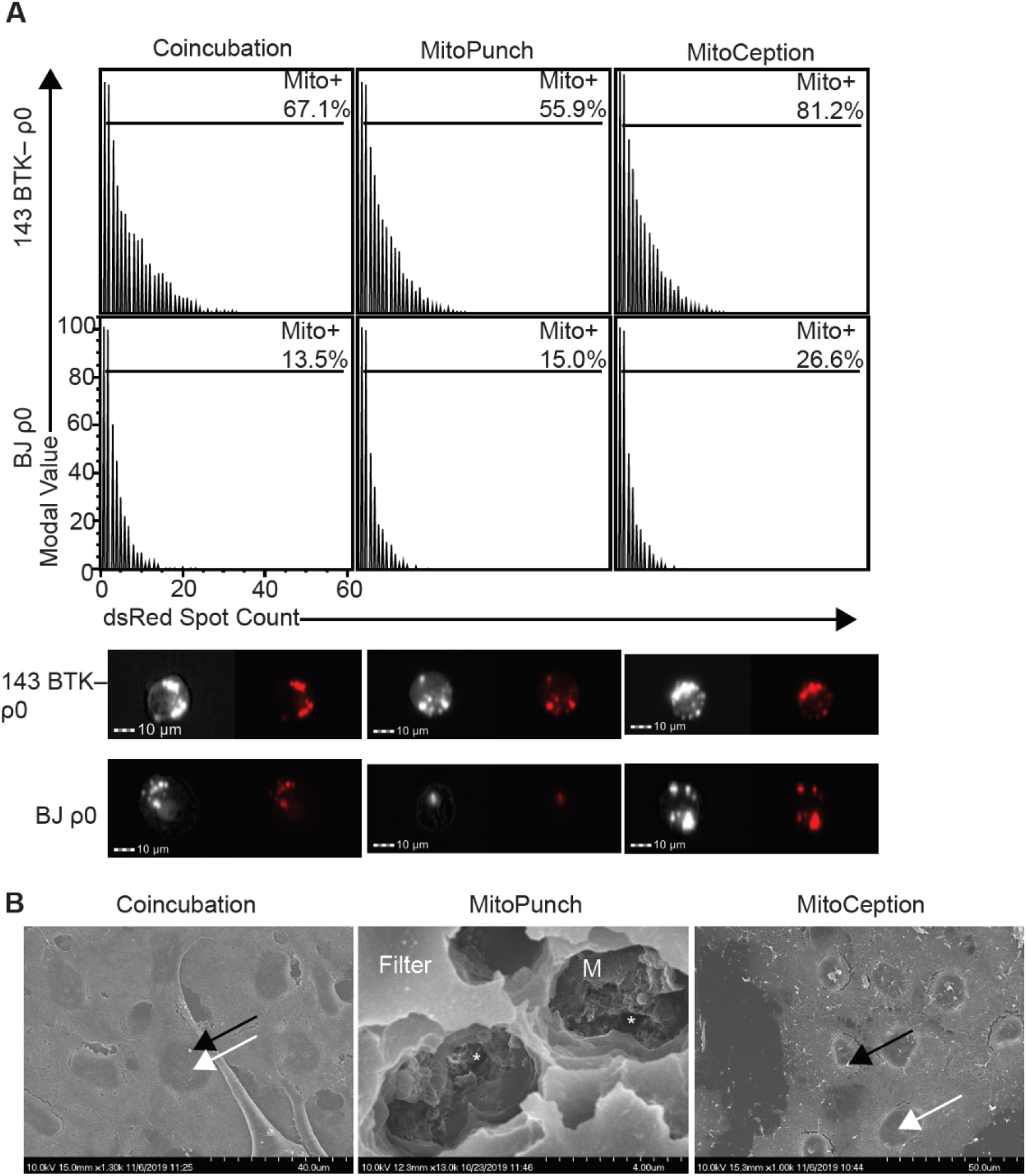
Mitochondrial spot quantification and SEM micrographs. **(A)** Representative bright-field and PE channel fluorescent images from ImageStream flow cytometry with spot count analysis of the number of mitochondria received by 143BTK− ρ0 and BJ ρ0 cells. Flow cytometry data is represented as histograms normalized to the mode. Scale bars indicate 10 μm. **(B)** SEM of 143BTK− ρ0 cells following mitochondrial transfer using the indicated techniques. Coincubation and MitoCeption images were taken from the apical face of the sample, whereas MitoPunch images were from the basal face of the sample through holes in the PET filter. In coincubation and MitoCeption images, the black arrows point to representative mitochondrial protein fragments and the white arrows point to representative cell nuclei. In the MitoPunch image, “Filter” indicates the PET filter membrane, “M” indicates the basal face of the cell membrane, and “*” indicates representative membrane damage.

**Supplemental Figure 2.**
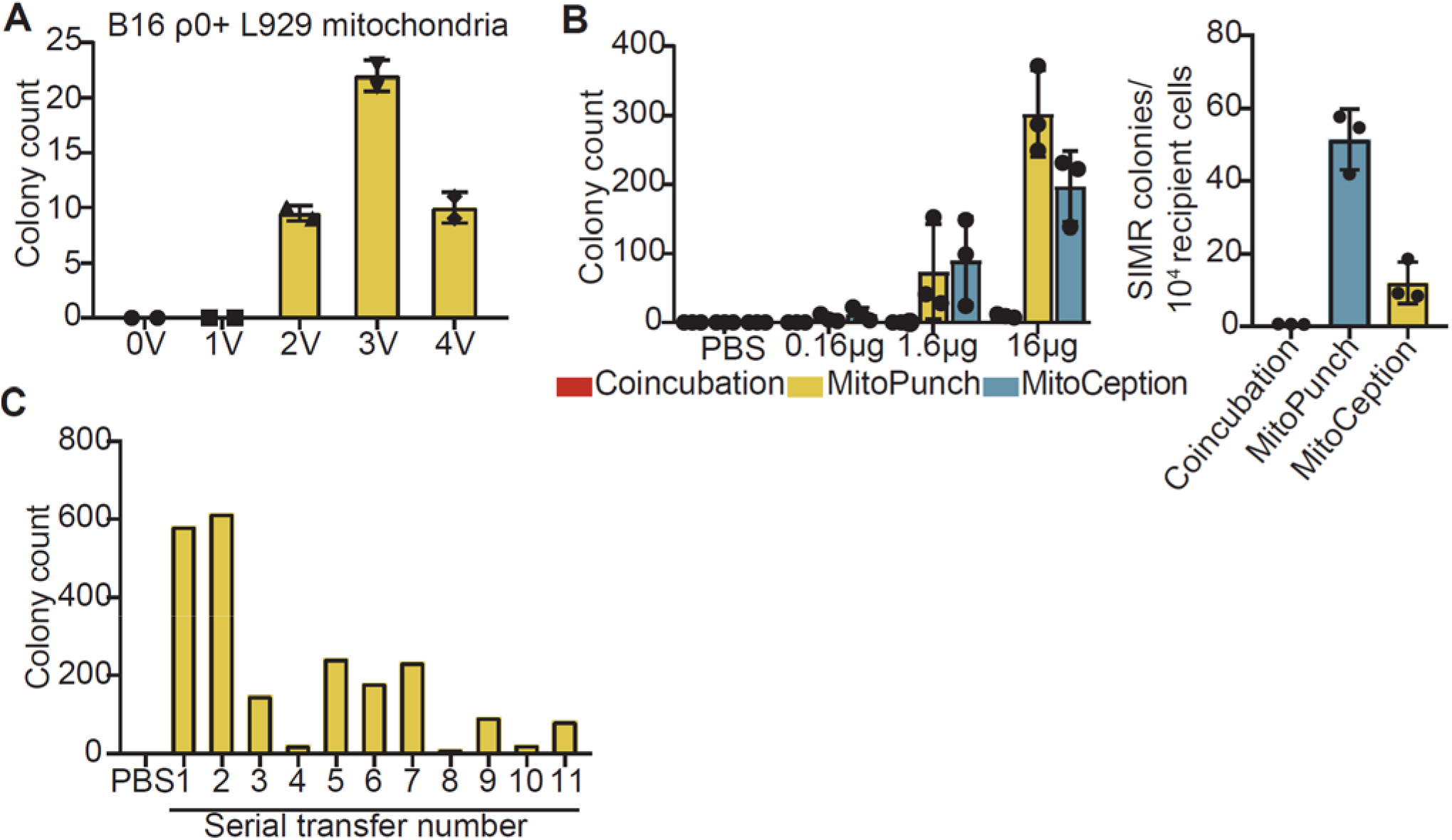
Quantification of stable mitochondrial delivery. **(A)** Quantification of crystal violet stained B16 ρ0 SIMR clones formed by MitoPunch actuated with indicated voltages after uridine-free selection. Error bars indicate the range of technical duplicates plotted with the average for each sample type. **(B)** Quantification of crystal violet stained 143BTK− ρ0 SIMR clones using indicated concentrations of mitochondrial suspension after uridine-free selection (left panel). 100% represents 16 μg mitochondria loaded into the PDMS delivery chamber. Error bars represent SD of three technical replicates. Quantification of SIMR clone generation efficiency for 1 x 104 cells that stained positive for dsRed protein by flow cytometry using 16 μg mitochondrial transfers (right panel). Error bars represent SD of three technical replicates. **(C)** Quantification of crystal violet stained 143BTK− ρ0 SIMR clones formed by serial MitoPunch deliveries of HEK293T mitochondria using the same sample remaining in the PDMS reservoir after the preceding delivery.

**Supplemental Figure 3.**
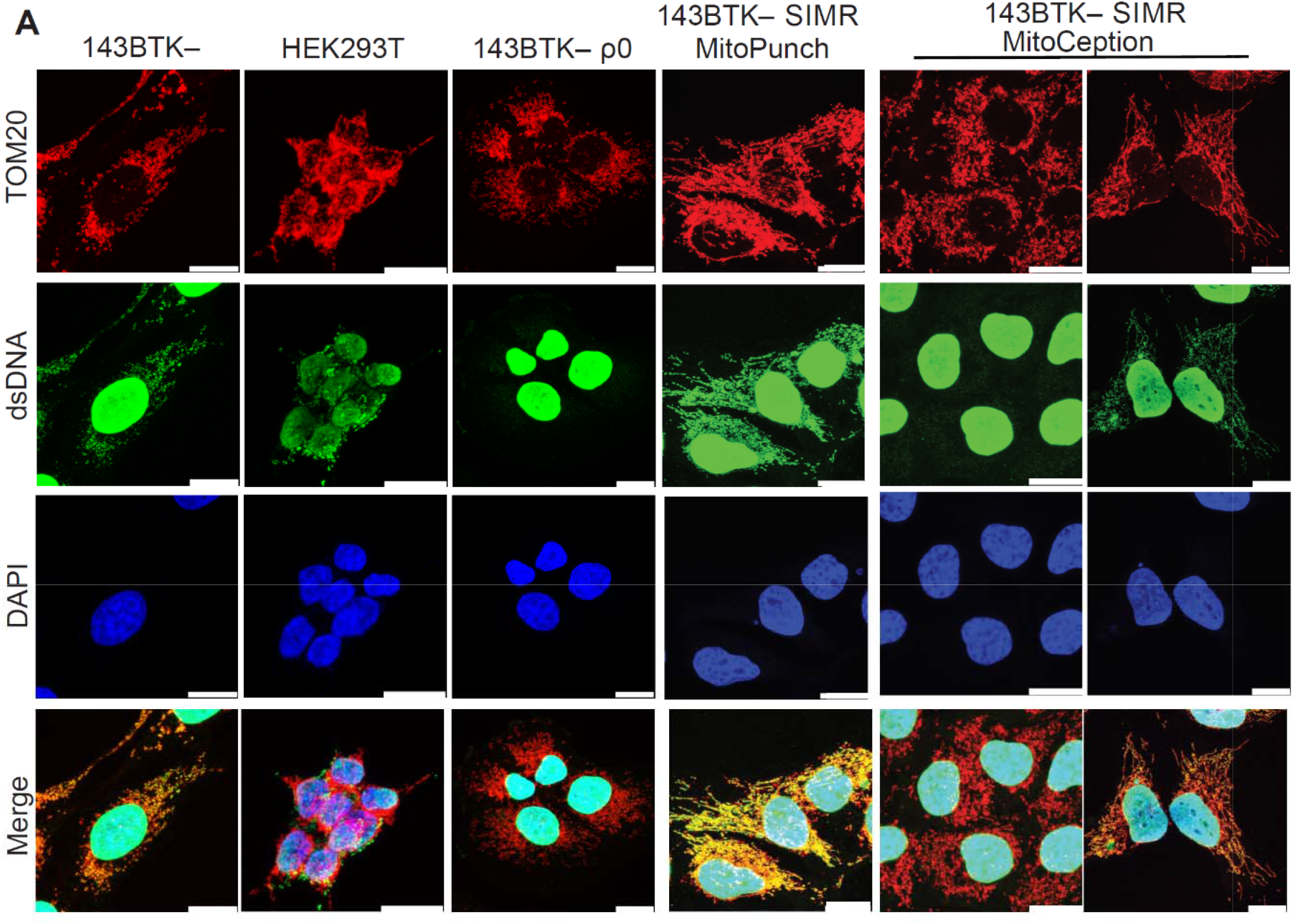
Confocal microscopy of stable mtDNA transplant lines. **(A)** Confocal microscopy of SIMR clones formed in 143BTK− ρ0 cells with 143BTK− parental, HEK293T dsRed mitochondrial donor, and 143BTK− ρ0 controls. The 143BTK-, 143BTK− ρ0, and HEK293T dsRed control images are the same images used in Figure 4B. Scale bars indicate 15 μm.

